# Arginine synthesis pathway and ASS1 play a critical role in mRNA translation reprogramming and ICI resistance in cutaneous melanoma

**DOI:** 10.64898/2026.03.17.712479

**Authors:** Emilie Chessel, Wassila Khatir, Ana Carolina B. Sant’Anna-Silva, Patricia Abbe, Guillaume Beranger, Zheyu Ding, Thierry Passeron, Issam Ben-Sahra, Henri Montaudié, Caroline Robert, Shensi Shen, Stephane Rocchi, Michaël Cerezo

## Abstract

Immune checkpoint inhibitors (ICI) have revolutionized cancer treatment, but their efficacy has now reached a plateau. ICIs are the first class of treatment targeting the crosstalk between immune and tumor cells, making it crucial to understand the complex interactions within the tumor microenvironment (TME) to enhance therapeutic responses. The elevated consumption of resources by cancer cells, coupled with limited vascularization, often results in a TME that is deficient in nutrients, leading to competition for resources between cancer and stromal cells. Consequently, targeting tumor metabolism has emerged as a promising strategy to improve the efficacy of ICIs. Through metabolomic analysis, we have identified metabolic alterations in melanoma cells that are resistant to ICIs, specifically an increase in arginine synthesis and upregulation of ASS1, the rate-limiting enzyme in this pathway.

By using gain and loss of function models, as well as a pharmacological inhibitor specific for ASS1, we demonstrated that modulations in the expression or activity of ASS1 is associated with translational reprogramming, characterized by an inhibition of the cap-dependent mRNA translation mediated through mTORC1/4EBP1 axis. We also demonstrated that targeting ASS1 *in vivo*, resensitize tumors initially resistant to ICI.

Taken together, our results highlight the interaction between modulations of arginine synthesis pathway, mRNA translation reprogramming, antitumor immunity, and restauration of sensitivity to anti-PD-1. Our work also demonstrates the therapeutic potential of targeting arginine synthesis pathway, and especially ASS1, to offer new treatments to patients suffering from cutaneous melanoma resistant to ICIs.

## Introduction

Over the past fifteen years, the paradigm of cancer therapy has been revolutionized using immune checkpoint inhibitors (ICI), in which the goal is to boost anti-tumor immunity in contrast with previously stablished therapies aiming at inducing cancer cells death. However, despite this breakthrough, in which cutaneous melanoma was leader, we have now reached a plateau. In cutaneous melanoma, ∼50% of patients do not respond to immunotherapy or become resistant to currently available strategies (Rohaan, Borch et al. 2022, Tawbi, Schadendorf et al. 2022, Wolchok, Chiarion-Sileni et al. 2025). Therefore, understanding the molecular mechanisms driving this resistance is essential to identify patients potentially susceptible to develop a good response and to propose new alternative strategies to treat resistant patients.

In this context of ICI usage, understanding the complex relationship taking place in the tumor microenvironment (TME) is crucial to improve responses. Importantly, due to the consumption of resources by cancer cells and poor vascularization, the TME frequently lacks nutrients and oxygen, which contributes to competition for nutrient access between cancer and stromal cells (Sukumar, Roychoudhuri et al. 2015, Reinfeld, Madden et al. 2021). In order to maintain their high proliferation rate and their survival in hypoxic and nutrients deprived microenvironment, malignant cells must adapt their metabolism (Icard and Lincet 2012, Hanahan 2022).

Among all the nutriments necessary for tumor cells, amino acids are particularly interesting for their central role in cell homeostasis and their role as elementary building blocks of polypeptide chains. Among all the amino acids, arginine is the most depleted one in the core of the tumors, leading to a competition between tumor cells and surrounding cells, such as CD8 positive T lymphocytes (Pan, Reid et al. 2016, Sullivan, Danai et al. 2019, Vecchio, Caiazza et al. 2021). In addition, arginine is semi-essential. It can be uptake from the TME or synthetized by the urea cycle. Aspartate is used to fuel the urea cycle and is catabolized by arginosuccinate synthetase (ASS1), the rate limiting enzyme of arginine synthesis that will combine it with citrulline to produce arginosuccinate, and argininosuccinate lyase (ASL) that will convert arginosuccinate into arginine (Husson, Brasse-Lagnel et al. 2003). Interestingly, lack of ASS1 expression in different cancers, such as most cutaneous melanoma or hepatocellular carcinoma (HCC) results in an arginine uptake dependency (Szlosarek, Klabatsa et al. 2006, Feun, Marini et al. 2012). Thus, the capacity of cancer cell to deal with low availability of arginine by overcoming their capacity to synthetize them, make some cancers more or less auxotrophic for arginine, opening a potential Achille heel that could be exploited to improve cancer treatments. In the last 20 years, different groups proposed a new therapeutical approach for melanoma (Ott, Carvajal et al. 2013) and HCC (Abou-Alfa, Qin et al. 2018) deficient for ASS1 by degrading arginine with arginine deiminase analogous ADIPEG20. However, despite an initial inhibition of tumor growth, tumor cells rapidly developed compensatory mechanisms by increasing ASS1 expression to maintain arginine synthesis (Tsai, Aiba et al. 2012, Long, Tsai et al. 2013). Thus, identifying alternative strategies to target the arginine pathway is necessary to exploit the therapeutic window offered by tumor cell dependency on arginine.

Arginine is with leucine and methionine, one of the three amino acids able to directly regulate mTORC1 complex though its sensors call CASTOR (Chantranupong, Scaria et al. 2016). mTORC1 complex is a central hub involved in tumor growth and proliferation by promoting anabolic functions such as protein synthesis (Morita, Gravel et al. 2015). The mTORC1/p70ribosomal S6 kinase axis regulates the ribosomes biogenesis, and the mTORC1/4EBP1 axis will allow the formation of the translation initiation complex eIF4F via the release of one of its subunits eIF4E after 4EBP1 phosphorylation (Laplante and Sabatini 2009, Laplante and Sabatini 2012). Thus, arginine appears to be at crossroads of tumor cell metabolism and mRNA translation, two major drivers of cancer aggressivity and therapy resistance (Fabbri, Chakraborty et al. 2021, Tang, Li et al. 2024).

Since targeting tumor metabolism is a keystone for development of new strategies to improve response to ICI and bypass resistance (Cerezo and Rocchi 2020), we investigated the metabolic adaptation associated with ICI resistance. In this study, we identify that arginine synthesis pathway and its limiting enzyme ASS1 are upregulated in cancer cell resistant to ICI, increasing the arginine availability, activating mTORC1 pathway and finally orchestrating a mRNA translation reprogramming driving resistance to anti-PD1 immunotherapy.

## Results

### ICI resistance is associated with arginine synthesis reprogramming

To dissect metabolic reprogramming associated with ICI resistance, we decided to use mouse melanoma cell lines. We first confirmed that the YUMM2.1 cells are sensitive to anti-PD-1 and YUMM1.1 cells are resistant. For this, we monitored tumor growth for 19 days and observed that anti-PD-1 significantly reduces tumor growth by 50% in YUMM2.1-injected mice, whereas YUMM1.1 derived tumors were not affected by this treatment (Figure 1A). Then, to study metabolic rewiring involved in the development of resistance we first performed a steady state metabolomic analysis comparing YUMM1.1 and YUMM2.1 cells and we observed a dysregulation at the crossroad between pyrimidine synthesis and urea cycle (Figure 1B). To follow specifically the urea cycle and characterize its potential dysregulations in resistant cells, we used ^15^N-amine-labeled *L*-glutamine. This approach allows us to identify an upregulation of arginine synthesis fuel by an upregulation of the glutaminolysis in arginine in ICI resistant cell line (YUMM1.1) compared to the sensitive one (YUMM2.1) (Figure 1C-D). We also performed this assay to follow the pyrimidines synthesis pathway; we observed as well an increase in this pathway between ICI-resistant YUMM1.1 and ICI-sensitive YUMM2.1 coherent we the increase of glutaminolysis (Suppl. Fig. 1). Then we monitored the expression level of ASL and ASS1 enzymes; the latter is the rate limiting enzyme for arginine synthesis. Consistent with the increase of arginine synthesis, we observed an increase of ASS1 and ASL both at the mRNA (Figure 1E) and protein levels (Figure 1F) in ICI-resistant cells YUMM1.1 compared to ICI-sensitive cells YUMM2.1. Finally, we analyzed two different cohorts of melanoma patients, and we observed an increase of ASS1 mRNA expression in tumors from patients that do not respond to ICI therapies (Figure 1G).

**Figure 1:**
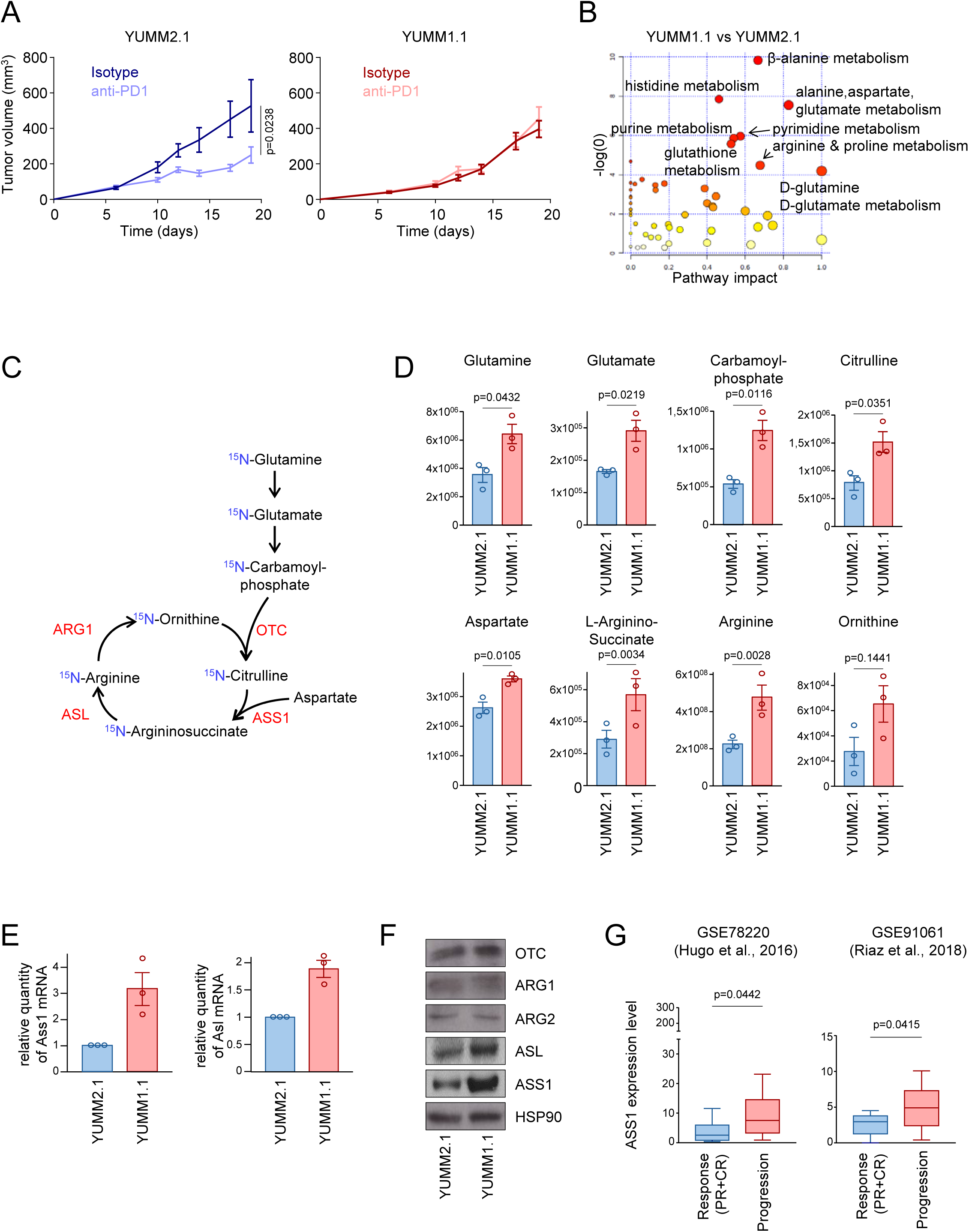
Resistance to ICI is associated with arginine synthesis reprogramming. **A**, Mice were engrafted with YUMM2.1 (left panel) or YUMM1.1 (right panel) and treated with anti-PD1. The tumor growth was followed for 3 weeks. **B**, Steady state metabolomic data from sensitive and resistant murine melanoma cells were analyzed with Metaboanalyst and represent using Pathway Impact package. **C**, Schematic representation of the urea cycle fueled from glutamine. **D**, Normalized peak areas of ^15^N-labeled metabolites measured by targeted LC-MS/MS from sensitive and resistant. **E**, RT-qPCR showing the expression level of the indicated mRNA. **F**, Immunoblot showing the indicated protein expression level in the YUMM2.1 and YUMM1.1 cells. **G**, Expression level of ASS1 mRNA in melanoma biopsies from patients that respond or not to ICI.

Altogether, our data demonstrate that melanoma cells resistant to ICI reprogram their arginine synthesis pathway by increasing mRNA and protein levels of ASS1 and ASL, to increase their intracellular levels of arginine.

### Cap-dependent mRNA translation is up regulated in melanoma cells resistant to ICI

To further explore the consequences of the increase of arginine synthesis in ICI-resistant cells, we performed a Gene Set Enrichment Analysis (GSEA), and we observed an enrichment of mRNA signatures involved in functions related to the formation of ribosomes and involved in the regulation of mRNA translation in YUMM1.1 ICI-resistant cells compared to YUMM2.1 ICI-sensitive cells (Figure 2A). To validate these differences in mRNA translation machinery, we compared expression levels of proteins involved in this mechanism by western-blot analysis. In the ICI resistant cells, we observed a decrease of 4EBP1 expression, a key protein that controls the formation of the translation initiation complex, associated with an increase in its phosphorylated forms. In contrast, we didn’t observe any differences in the expression levels of the 3 sub-units, eIF4A, eIF4E and eIF4G, forming the translation initiation complex eIF4F (Figure 2B). To explore the modulations of translation in ICI resistant cell lines, we first transfected YUMM2.1 and YUMM1.1 cells with a plasmid containing two coding sequences for a cap-dependent Renilla luciferase and a cap-independent Firefly luciferase linked through an IRES allowing to distinguish the cap-dependent from the cap-independent translation respectively (Figure 2C, upper panel). We observed an increase in the Renilla over Firefly luminescent ratio in the YUMM1.1 cells reflecting the increase of the ratio CAP-dependent to CAP-independent translation (Figure 2C, lower panel).

**Figure 2:**
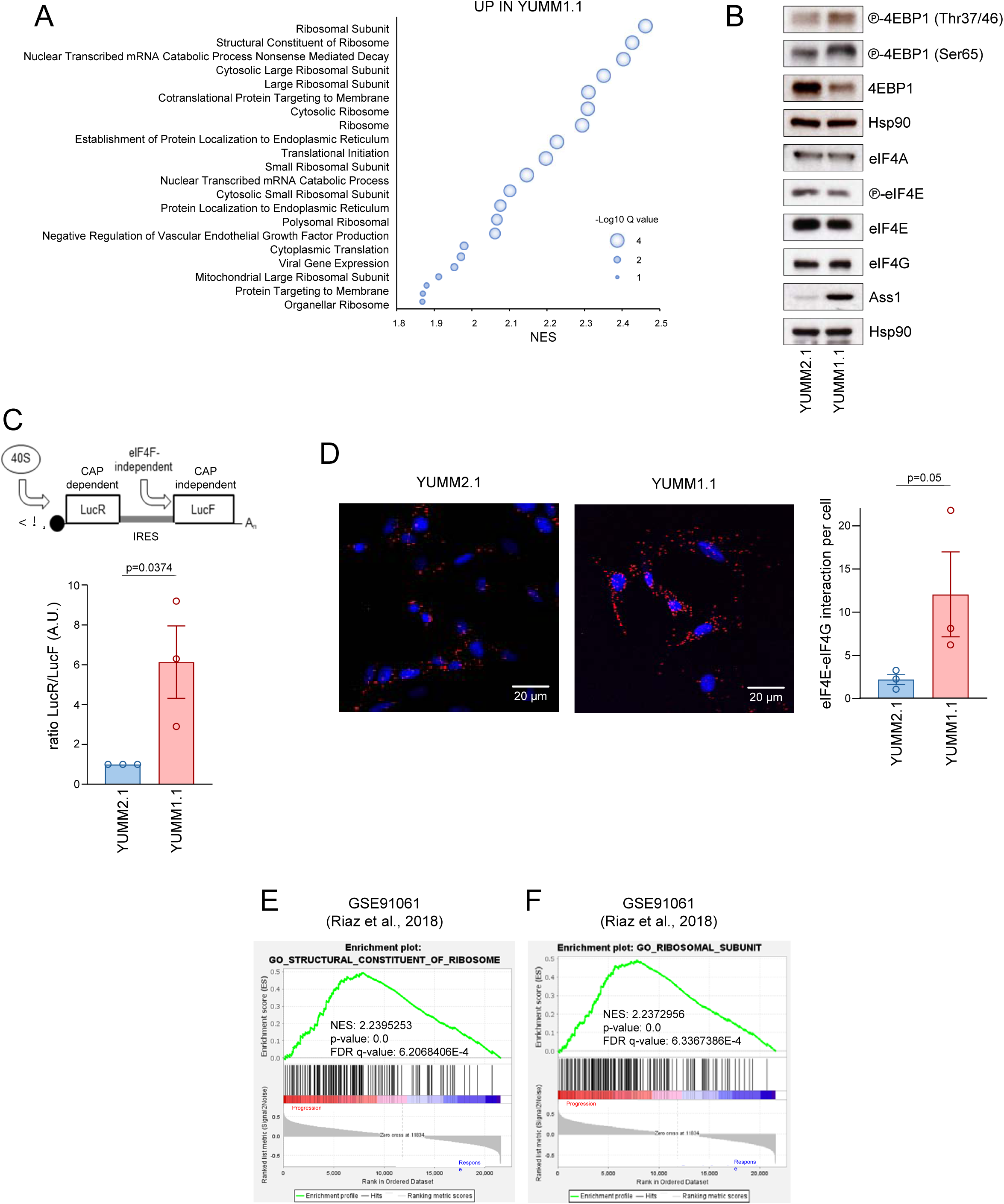
Cap-dependent mRNA translation is up regulated in melanoma cells resistant to ICI. **A**, GSEA analysis showing a set of genes significantly dysregulated in ICI-resistant cells (YUMM1.1) compared to ICI sensitive cells (YUMM2.1). **B**, Western blot showing indicated proteins expression levels in ICI-sensitive YUMM2.1 cells versus ICI-resistant YUMM1.1 cells. HSP90 serves as a loading control. **C**, Representation of the bicistronic plasmid used to compare cap-dependent and independent translation initiation (upper panel). Renilla over Firefly luminescent ratio quantification in ICI-sensitive YUMM2.1 cells versus ICI-resistant YUMM1.1 cells (n=3; mean ± SEM) (lower panel). **D**, Proximity Ligation Assay showing eiF4E/eiF4G interactions in YUMM2.1 and YUMM1.1 cells. Representative images are presented in the left panel, and quantifications are presented in the right panel. **E-F**. Representation of enriched sets of genes, obtained after GSEA analysis, in patients that progress under ICI treatment compared to patients that respond.

We then performed a proximity ligation assay between eIF4E and eIF4G, two major components of eIF4F complex, to monitor their association. We observed an increase of the association between eIF4E and eIF4G and consequently an activation of the translation initiation complex in the YUMM1.1 resistant cells, compared to YUMM2.1 sensitive cells (Figure 2D). Finally, we performed a GSEA analysis in two datasets from patients responding or not to ICI and we observed an enrichment of signatures associated to mRNA translation in melanoma patients that progress under ICI treatment compared to patients that respond to the therapy (Figure 2E and 2F).

Together, our data demonstrate that mRNA translation pathways, and especially translation initiation, are upregulated in melanoma cells resistant to ICI.

### ASS1 controls mRNA translation in ICI-R melanoma cell lines

To further dissect the effect of ASS1 and consequently arginine pathway modulations on translational modifications, we used two different murine melanoma cell lines resistant to ICI (YUMM1.1 and YUMM1.7) and one murine melanoma cell line sensitive to ICI (YUMM2.1) (Meeth, Wang et al. 2016). To address the role of ASS1 on the control of translation, we used viral particles containing either a shRNA targeting ASS1 or a non-regulatable form of ASS1 under the control of a strong promoter to infect ICI-resistant and ICI-sensitive cells respectively. We first performed arginine uptake assay to verify that ASS1 inhibition was not compensate by an upregulation of the intake (Suppl. Fig. 2). Then, we observed a decrease in the phosphorylation of 4EBP1 in both ICI-resistant cell lines when ASS1 is inhibited (Figure 3A, 3B). At the contrary, the overexpression of ASS1 leads to an increase of 4EBP1 phosphorylation (Figure 3C). As mTORC1 pathway is a major regulator of cell size we observed a correlation between the expression of ASS1 and the cell size (Suppl. Fig. 3). To go further in the role of ASS1 in the translation regulation, we transfected the cells with the plasmid previously described in Figure 2D, allowing us to distinguish the cap-dependent from the cap-independent translation. We observed that the ASS1 inhibition led to a decrease of the ratio CAP-dependent to CAP-independent translation (Figure 3D-E) while the over-expression of ASS1 lead to an increase of the ratio CAP-dependent to CAP-independent translation (Figure 3F). Then we performed a proximity ligation assay between eIF4E and eIF4G to monitor the translation initiation complex assembly. Inhibition of ASS1 leads to a decrease of eIF4F assembly (Figures 3G and 3H). Finally, to monitor the global translational activity, cells were subjected to ^35^S-labeled methionine, to detect and quantify *de novo* protein synthesis. We observed a significant decrease in labeled methionine incorporation for both YUMM1.1 and YUMM1.7 expressing the shASS1 (Figure 3I, 3J), and a non-significant tendency to an increase in the labeled methionine incorporation in YUMM2.1 cells overexpressing ASS1 (Figure 3K).

**Figure 3:**
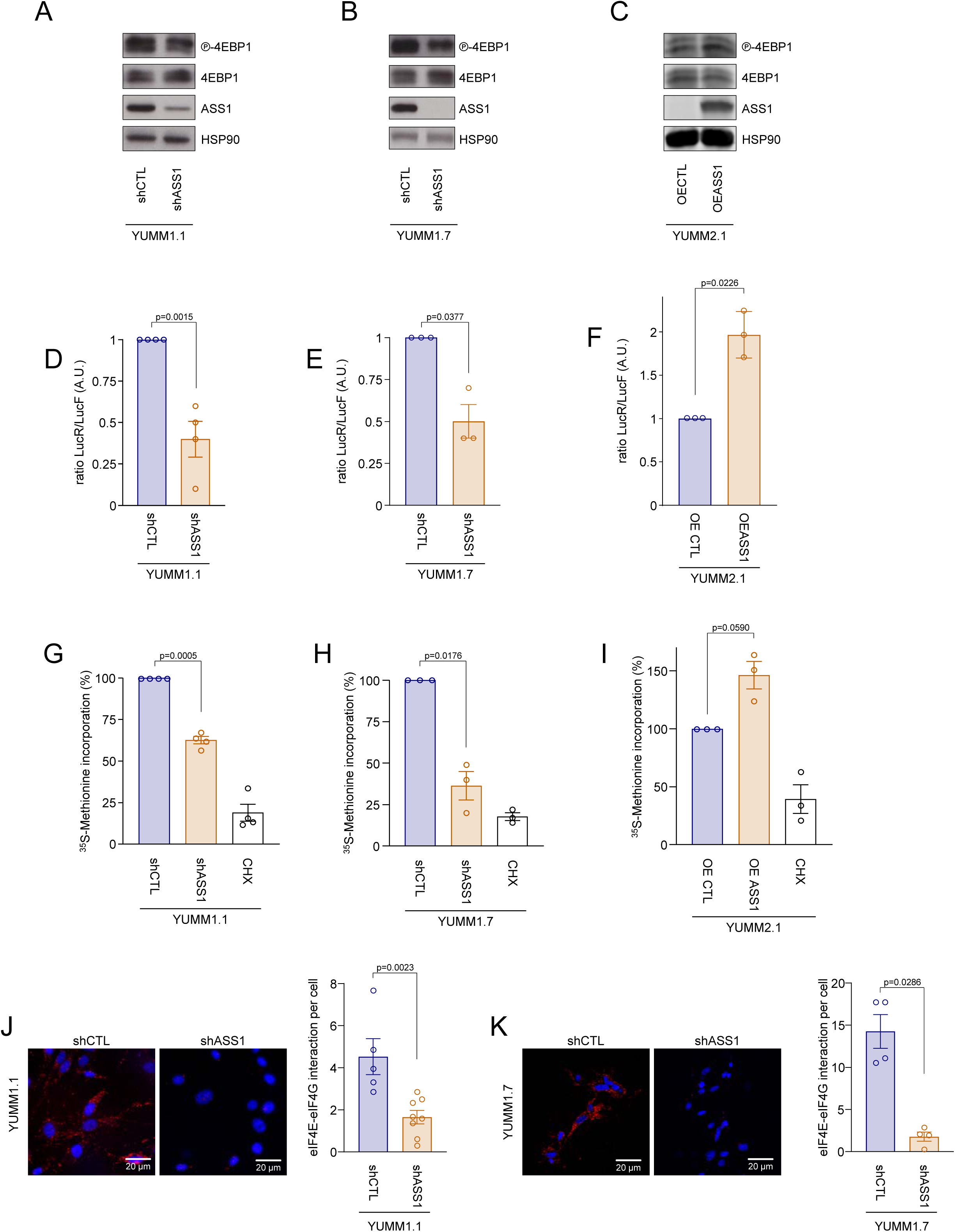
ASS1 controls mRNA translation in ICI-R melanoma cell lines. **A**, Western blot showing indicated proteins expression levels in YUMM1.1 cells expressing a shRNA control or targeting ASS1. HSP90 serves as a loading control. **B**, Western blot showing indicated proteins expression levels in YUMM1.7 cells expressing a shRNA control or targeting ASS1. HSP90 serves as a loading control. **C**, Western blot showing indicated proteins expression levels in YUMM2.1 cells expressing an ectopic empty vector or containing the open reading frame of ASS1. HSP90 serves as a loading control. **D**, Renilla over Firefly luminescent ratio quantification in YUMM1.1 cells expressing a shRNA control or targeting ASS1 (n=4; mean ± SEM). **E**, Renilla over Firefly luminescent ratio quantification in YUMM1.7 cells expressing a shRNA control or targeting ASS1 (n=3; mean ± SEM). **F**, Renilla over Firefly luminescent ratio quantification in YUMM2.1 cells expressing an ectopic empty vector or containing the open reading frame of ASS1 (n=3; mean ± SEM). **G**, Protein synthesis rates were determined in YUMM1.1 cells expressing a shRNA control or targeting ASS1. Cells were then pulsed for 30 min with [^35^S]Cys/Met, and the incorporation of ^35^S into proteins was quantified and normalized to the total protein amount. Cells treated 30 minutes with 100µg/mL of cycloheximide (CHX) are used as a positive control. The data are presented as the mean (n = 4). **H**, Protein synthesis rates were determined in YUMM1.7 cells expressing a shRNA control or targeting ASS1. Cells were then pulsed for 30 min with [^35^S]Cys/Met, and the incorporation of ^35^S into proteins was quantified and normalized to the total protein amount. Cells treated 30 minutes with 100µg/mL of cycloheximide (CHX) are used as a positive control. The data are presented as the mean (n = 3). **I**, Protein synthesis rates were determined in YUMM2.1 cells expressing an ectopic empty vector or containing the open reading frame of ASS1. Cells were then pulsed for 30 min with [^35^S]Cys/Met, and the incorporation of ^35^S into proteins was quantified and normalized to the total protein amount. Cells treated 30 minutes with 100µg/mL of cycloheximide (CHX) are used as a positive control. The data are presented as the mean (n = 3). **J**, Proximity Ligation Assay showing eiF4E/eiF4G interactions in YUMM1.1 cells expressing a shRNA control or targeting ASS1. Representative images are presented in the left panel, and quantifications are presented in the right panel. **K**, Proximity Ligation Assay showing eiF4E/eiF4G interactions in YUMM1.7 cells expressing a shRNA control or targeting ASS1. Representative images are presented in the left panel, and quantifications are presented in the right panel.

Taken all together, these results demonstrate that ASS1 exerts control on mRNA translation through the modulation of eIF4F.

### ASS1 controls anti-tumor immunity through mRNA translation reprogramming

To identify the specific subset of mRNAs whose translation may be regulated by ASS1, we performed polysome fractionation followed by RNA sequencing. Briefly, this technique consists in the separation of mRNA on a sucrose gradient based on their weight and consequently the number of ribosomes associated with them. By comparing ICI-resistant YUMM1.1 cells with ASS1 knockdown to YUMM1.1 cells expressing a control shRNA, we observed a slight decrease in the heaviest polysome fractions in the latter (Figure 4A), which is consistent with the previously observed reduction in methionine incorporation and eIF4F complex assembly (Figure 4A). Then the RNA sequencing on the heaviest fractions and total mRNAs were analyzed to calculate a translation efficiency (for each modulated mRNA, we calculated the ratio of counts in the heavy polysome fractions to the total mRNA), followed by a differential analysis using EdgeR. Even though we observed, as expected, down-regulated mRNAs in YUMM1.1 cells lacking ASS1, we also observed sets of mRNAs up-regulated when ASS1 is inhibited (Figure 4B and 4C). We then performed a GSEA analysis in up-regulated mRNAs and found out that the most enriched pathways are involved in antigen processing and presentation (Figure 4D). For example, among the upregulated mRNAs, we identified several transcripts involved in immunoproteasome formation, such as *Psmb9* and *Ubd*, as well as *H2-Q7*, *H2-Q10*, and *H2-M3*, which are involved in major histocompatibility complex (MHC) formation (Figure 4C).

**Figure 4:**
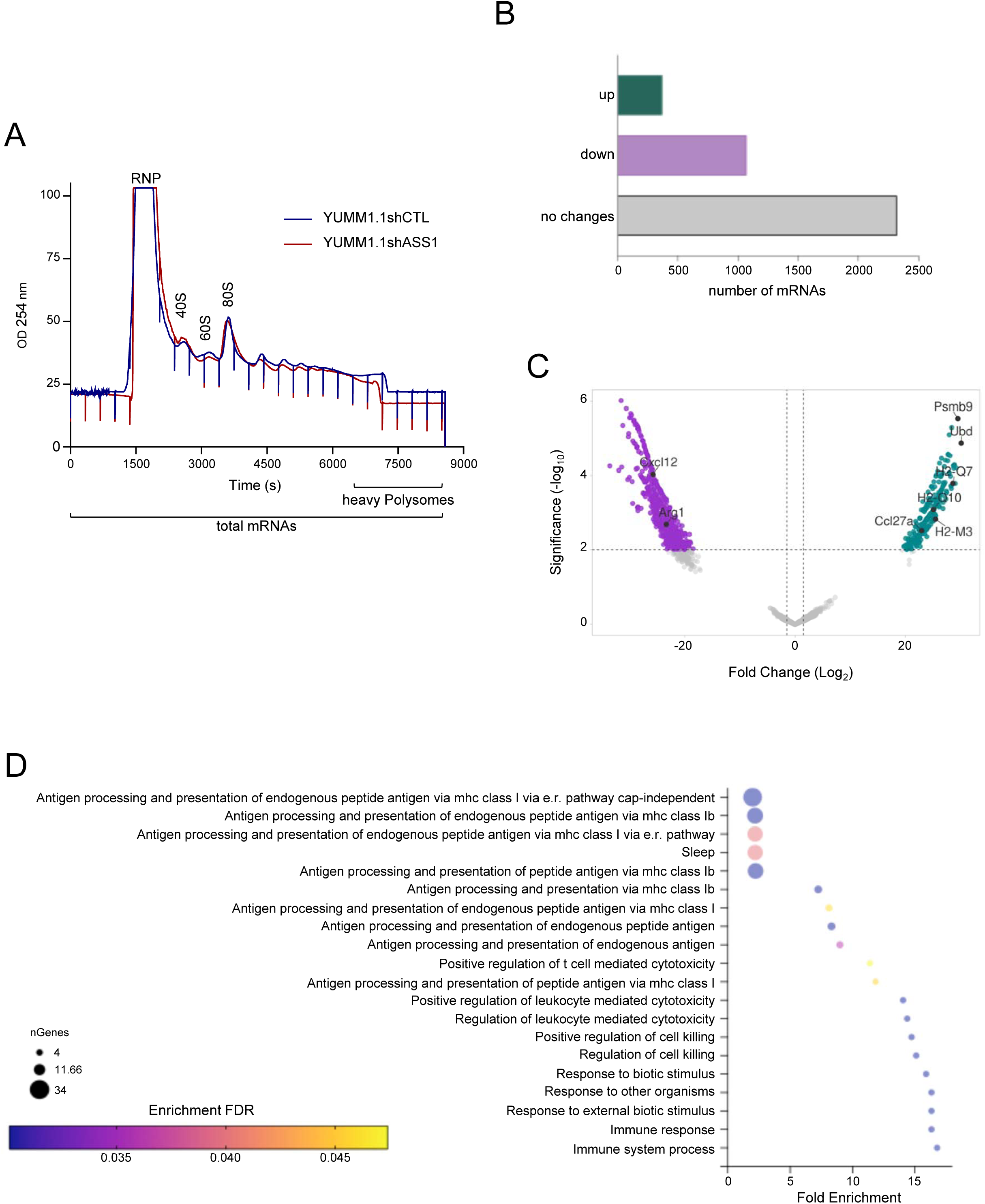
ASS1 controls anti-tumor immunity and ICI-R through mRNA translation. **A**, Polysome profiles of YUMM1.1 cells expressing a shRNA control or targeting ASS1. OD, optical density; RNP, ribonucleoparticle. **B**, Numbers of mRNAS up-regulated (log2(fold change>1.5), down-regulated (log2(fold change<-1.5) or unchanged, in ICI-resistant YUMM1.1 cells knocked-down for ASS1 compared to ICI-resistant YUMM1.1 expressing a shRNA control. Analysis was performed with edgeR software, on the data of RNA sequencing of the highest fractions of polysomes (heavy polysomes highlighted in Figure 4A). **C**, Volcano plot shows mRNAs of immunoproteasome and MHC class I and class II proteins whose expression is increased (log2(fold change)>20, p-values <0,005) (green dots) or decreased (log2(fold change)<-20, p-values <0,005) (purple dots) upon highest polysomes fractions. VolcanoNoseR was used to obtain the representation. **D**, GSEA analysis showing pathways that are significantly up regulated in the highest polysomes fractions.

These data suggested that that ASS1 inhibition in ICI-resistant cells might, by inhibiting CAP-dependent translation and consequently liberate a pool of ribosomes, promote the translation of a subset of mRNA implicated in anti-tumor immunity. Thus, these data suggest the implication of ASS1 in the translational control of anti-tumor immunity.

### Modulations of ASS1 restore anti-PD-1 sensitivity *in vivo*

Given the concordance between arginine synthesis modulations, translational reprogramming, and ICI-resistance, we explored the effect of ASS1 modulations on the response to anti-PD-1 treatment *in vivo*. We used immunocompetent C57BL/6 mice in which we engrafted YUMM1.1 resistant cells expressing or not a shRNA targeting ASS1 (Figure 5A). We followed the tumor growth for 18 days, and we observed, at a dose of 4mg.kg^-1^ of anti-PD-1, a significant inhibition of tumor volume in ICI resistant cells YUMM1.1 inhibited for ASS1 (Figures 5A and 5B). To characterize the impact of ASS1 modulations on anti-tumor immunity, we performed flow cytometry experiments on dissociated tumors, and we observed an increased amount of CD45^+^ cells in tumors derived from ASS1 *knocked-down* cells and treated with anti-PD-1 (Figure 5C). To go further, we performed immunofluorescence assay on tumors sections, and we observed a significant increase of the CD8^+^ cells infiltration in tumors derived from YUMM1.1 cells *knocked-down* for ASS1 and treated with anti-PD-1 (Figures 5D, 5E). To evaluate the CD8^+^ T cells activity in the tumors, we performed granzyme B immunofluorescence assay (Cullen, Brunet et al. 2010). We didn’t observe granzyme B secretion in tumors derived from YUMM1.1 expressing a shRNA control treated with anti-PD-1 reflecting the probable state of exhaustion of the CD8 positive cells presented in these tumors. However, we observed a strong increase of the granzyme B staining in tumors lacking ASS1 and treated with anti-PD-1 (Figure 5F) demonstrating that the inhibition of ASS1 combine with anti-PD1 treatment can stimulate an infiltration of activated anti-tumor CD8 positive cells within the tumors.

**Figure 5:**
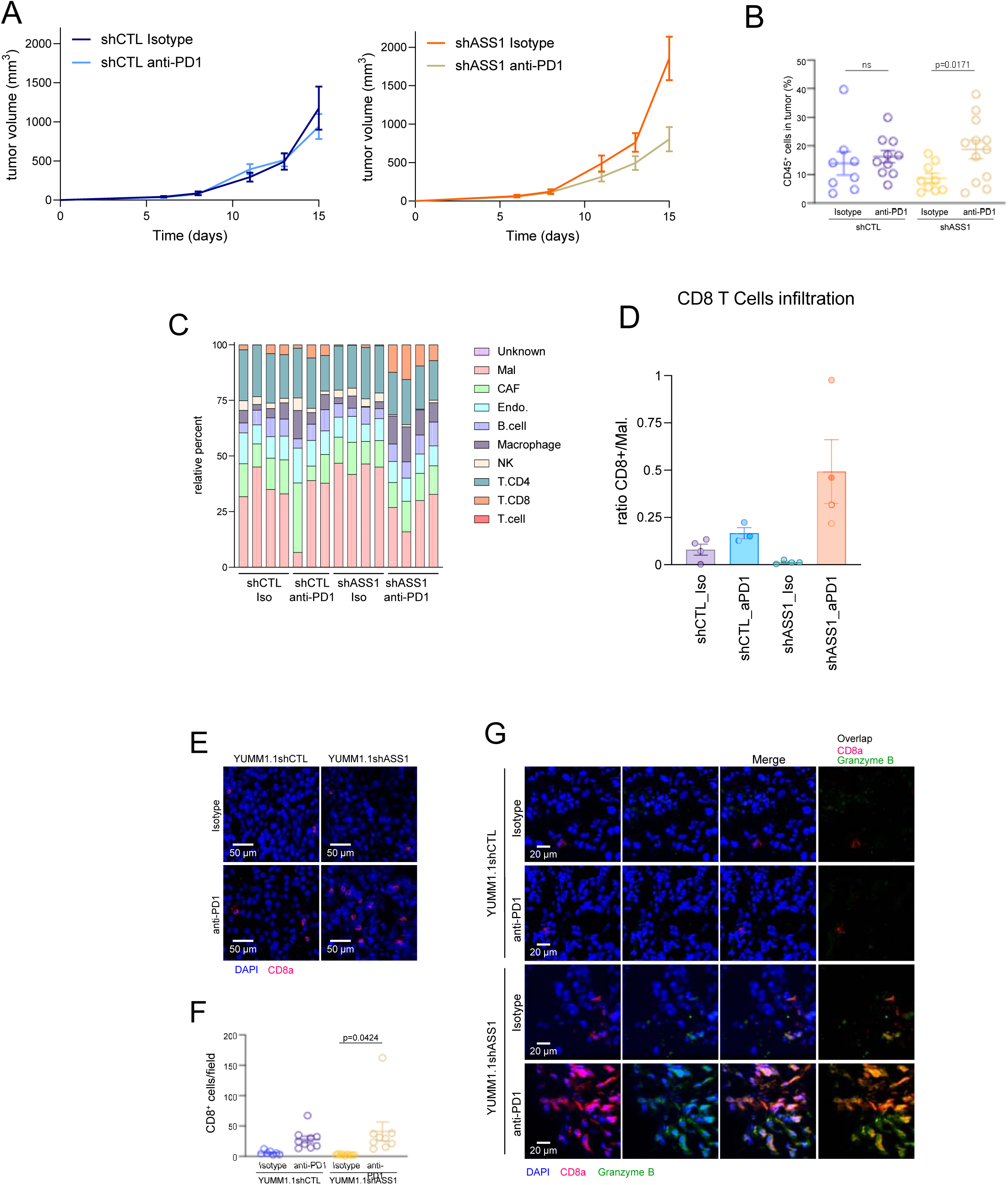
ASS1 modulates anti-tumor immunity by reshaping tumor infiltrate. **A**, C57/BL6 mice were inoculated with murine Braf^V600E^ Pten^-/-^ melanoma cells (YUMM1.1), expressing a shRNA targeting ASS1 or a shRNA control. When tumors reached a mean of 100mm3, mice were treated with anti-PD-1 (4mg/kg-1) or isotype IgG control. The data are presented as the mean ± SEM for each group (n=10 mice per group). p-values were calculated using two-way ANOVA. **B**, Percentage of CD45-positive cells in tumors was determined by flow cytometry in tumors presented in B. The data are presented as the mean ± SEM (YUMM1.1shCTL n=9 tumors ; YUMM1.1shASS1 n=10 tumors). **C**, Deconvolution performed using CIBERSORTx performed on tumors from the experiment presented in A. **D**, Ratio CD8+ cells to malignant cells (Mal.) from the panel C is presented in tumor from the experiment presented in the panel A. **E**, Representative immunostaining of CD8+ T lymphocytes (red, CD8a) and nuclei (blue, DAPI) on frozen tumor sections of mice from panel A. **F,** Representation of CD8+ cells shown in D, the data are presented as the mean ± SEM (n=9 tumors). **G.** Representative immunostaining of CD8+ T cells and granzyme B on frozen tumor sections of mice from panel A.

Altogether, our results demonstrate that genetic inhibition of ASS1 resensitize tumors to anti-PD-1 treatment by stimulating the CD8+ immune cells infiltration and their activation within the tumors.

### Pharmacological inhibition of ASS1 activity inhibits CAP-dependent translation and restore anti-PD-1 sensitivity *in vivo*

To confirm the results obtained with the genetic inhibition of ASS1, we used the lZl-methyl-*DL*-aspartic acid (MDLA), an analog of aspartate that inhibits specifically ASS1 enzymatic activity. MDLA treatment did not affect ASS1 or ASL expression levels in ICI-resistant YUMM1.1 cells (Figure 6A). However, we observed a significant decrease of ASS1 activity in a dose-dependent manner in ICI-resistant cells treated with MDLA (Figure 6B), associated with a significant decrease in intracellular arginine concentration at the highest dose of MDLA (Figure 6C).

**Figure 6:**
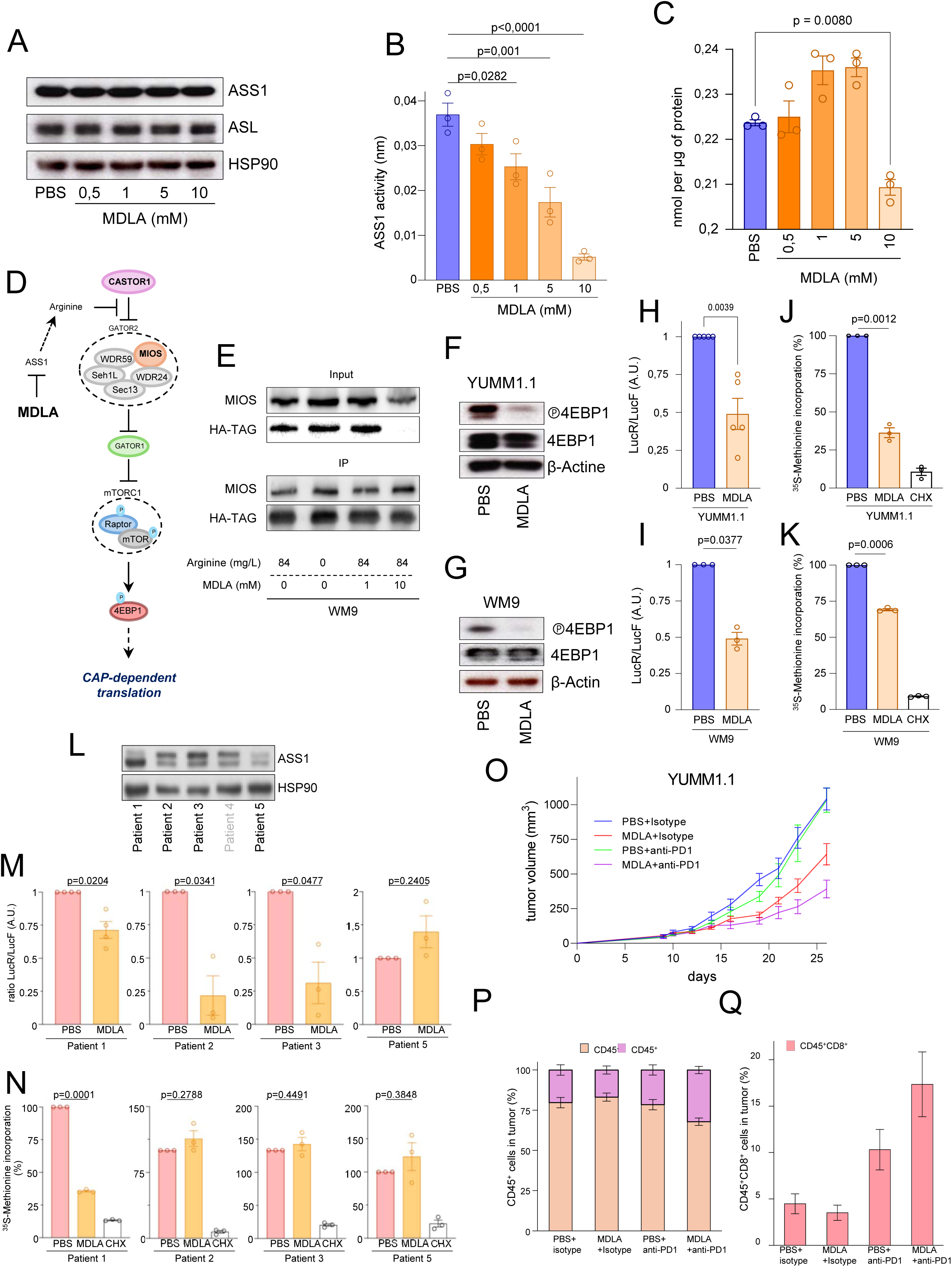
Targeting of ASS1 revert ICI resistance. **A**, Immunoblot showing the expression levels of indicated protein in ICI-resistant YUMM1.1 cell line after 3 days of treatment with MDLA. HSP90 serves as a loading control. **B**, ASS1 enzymatic activity was performed in ICI-resistant YUMM1.1 cell line after 3 days of treatment with (n=3; mean ± SEM). **C**, Intracellular concentration of arginine was measure in ICI-resistant YUMM1.1 cells after 3 days of treatment with MDLA. (n=3; mean ± SEM). **D.** Schematic representation of the impact of arginine on the control upstream of mTORC1. **E.** Immunoprecipaption was performed on WM9 cells transfected with HA-tagged CASTOR1 and cultured for 3 days in the presence of different concentration of arginine and MDLA. Immunoblot showing the expression levels of indicated protein. **F**, Immunoblot showing the expression levels of indicated protein in ICI-resistant YUMM1.1 cell line. β-actine was use as a loading control. **G**, Immunoblot showing the expression levels of indicated protein in WM9 melanoma cell line. **H**, Renilla over Firefly luminescent ratio quantification in ICI-resistant YUMM1.1 cell line. **I**, Renilla over Firefly luminescent ratio quantification in WM9 melanoma cell line. **J**, YUMM1.1 cells treated or not with MDLA were then pulsed for 30 min with [^35^S]Cys/Met, and the incorporation of ^35^S into proteins was quantified and normalized to the total protein amount. Cells treated 30 minutes with 100µg/mL of cycloheximide (CHX) are used as a positive control. The data are presented as the mean (n = 3). **K**, WM9 cells treated or not with MDLA were then pulsed for 30 min with [^35^S]Cys/Met, and the incorporation of ^35^S into proteins was quantified and normalized to the total protein amount. Cells treated 30 minutes with 100µg/mL of cycloheximide (CHX) are used as a positive control. The data are presented as the mean (n = 3). **L**, Immunoblot showing the expression levels of indicated protein in cells from tumors of patients with metastatic melanoma. HSP90 serves as a loading control. **M**, Renilla over Firefly luminescent ratio quantification in cells from tumors of patients with metastatic melanoma (n=3; mean ± SEM). **N**, Cells isolated from tumors of patients with metastatic melanoma were treated or not with MDLA were then pulsed for 30 min with [^35^S]Cys/Met, and the incorporation of ^35^S into proteins was quantified and normalized to the total protein amount. Cells treated 30 minutes with 100µg/mL of cycloheximide (CHX) are used as a positive control. The data are presented as the mean (n = 3). **O**, C57BL/6 mice were inoculated with murine *Braf^V600E^ Pten^-/-^* melanoma cells (YUMM1.1). When tumors reached a mean of 100mm^3^, mice were treated with MDLA (700mg.kg-1) or PBS, twice a day. Two days after the beginning of the MDLA treatment, mice were treated with anti-PD-1 (4mg/kg-1) or isotype IgG control, once per day, every two days, 3 times. Mice were treated with an additional dose of anti-PD-1 or isotype IgG control at day 24. Tumor growth was monitored for 27 days. The data representing the tumor growth are presented as the mean ± SEM for each group (n=10 mice per group). *p*-values were calculated using two-way ANOVA. **P**, Percentage of CD45-positive cells in tumors was determined by flow cytometry in tumors presented in P. The data are presented as a ratio of percentage between CD45-positive cells versus CD45-negative cells. (n=10 tumors per group, mean ± SEM). **Q**, Percentage of CD45 and CD8a-double positive cells in tumors, determined by flow cytometry in tumors presented in P. (n=10 tumors per group, mean ± SEM).

Given the impact of ASS1 modulations on mTORC1/4EBP1 axis, we explored the potential impact of the inhibition of ASS1 activity on CASTOR1-GATOR2 interaction (Chantranupong, Scaria et al. 2016) (Figure 6D). To do so, we expressed an HA-tagged form of CASTOR1 in WM9 melanoma cell line to monitor its interaction with GATOR2 complex. Interestingly, we observed that CASTOR1 co-immunoprecipitates with MIOS, an established GATOR2 component, in cells either treated with MDLA or depleted for arginine (Figure 6E). This result demonstrates that the pharmacological inhibition of ASS1 is sufficient to reduce arginine availability and drive an inhibition of mTORC1 complex. Then we explore the downstream effects of ASS1 activity blockage on mTORC1 pathway. We observed a decrease of the phosphorylated form of 4EBP1 in YUMM1.1 and WM9 treated with MDLA (Figure 6F and 6G). To study the impact of ASS1 activity blockage on mRNA translation, we transfected the cells with the plasmid previously described in Figure 2D and we observed a decrease of the Renilla over Firefly ratio in cells treated with MDLA reflecting an inhibition of CAP-dependent translation (Figure 6H and 6I). Then, we performed ^35^S-labeled methionine incorporation assay and we observed a significant decrease of the incorporation of labeled methionine in cells treated with MDLA (Figure 6J and 6K).

To confirm the effect of MDLA in a more physiopathological context, we used patient derived melanoma cells extracted from fresh biopsies of patients. We first measured the level of ASS1, and we found out a heterogeneity of protein expression depending on the patient (Figure 6L). Taking advantage of this observation, we chose 4 human-derived cell lines with different levels of ASS1 to evaluate the impact of ASS1 levels on the translational modulations induced by MDLA. After MDLA treatment, we observed a significant decrease in the CAP-dependent translation for the 3 patients with medium or high ASS1 expression (Figure 6M). In addition, we also observed a significant decrease in the ^35^S-methionine incorporation after MDLA treatment, only in cells with the highest ASS1 expression (Figure 6N).

We then explored to what extent MDLA could exert an immunotherapeutic effect *in vivo*. After engrafting immunocompetent C57BL/6 mice with ICI-resistant YUMM1.1, we treated them with MDLA (700mg.kg^-1^) or PBS twice a day for 16 days, associated with anti-PD-1 or isotype treatment. We followed the tumor growth for 27 days and observed a significant decrease in the tumor volume in MDLA+anti-PD-1 treated mice (Figure 6O). Finally, we explored the effect of MDLA might change the immune infiltrate in tumors, and we observed an increase of CD45^+^ cells in MDLA+anti-PD-1 treated mice (Figure 6P). Moreover, we observed an increase of CD45^+^/CD8^+^ T cells infiltrate in tumors treated with anti-PD-1 plus MDLA (Figure 6Q).

Taken together, our data demonstrate that targeting ASS1 activity with MDLA treatment leads to an inhibition of the CAP-dependent translation through a control of the mTORC1/4EBP1 axis. Moreover, MDLA treatment resensitize tumors to anti-PD-1 by stimulated CD8 driven anti-tumor infiltrate.

## Discussion

Our data show that the arginine synthesis pathway and more specifically ASS1 participate to resistance to ICI and especially anti-PD1 treatment through the control of a translation program implicated in the regulation of anti-tumor immunity.

Our data strongly suggest that cancer cell plasticity is a keystone for treatment resistance. Indeed, we are undercovering for the first time, a dialogue between *de novo* arginine synthesis pathway and mRNA translation supporting resistance to ICI. The coordination of metabolic and translational reprogramming has recently emerged as a new hub coordinating the strategies allowing cancer cells to escape from treatment (Tang, Li et al. 2024). Our study put the spotlight on the arginine pathway. Since decades, metabolic reprogramming has been associated with tumor aggressivity. The urea cycle and consequently arginine synthesis pathway has been described to be downregulated to fuel pyrimidine synthesis with aspartate supporting the high proliferation rate of cancer cells (Rabinovich, Adler et al. 2015, Gaude and Frezza 2016). However, we know now that the reality is much more complex and that ASS1 can be also upregulated in many cancers to supply cellular needs in arginine and downstream metabolites (Szlosarek, Grimshaw et al. 2007, Zhang, Liu et al. 2025). This relationship *between de novo* pyrimidine and arginine synthesis pathway has opened new therapeutic windows in the context of ICI. Indeed, a study have brilliantly demonstrated that high ASS1-expressing breast cancer mice do not respond to ICI, but that inhibiting purine pathway can in this context destabilized pyrimidine to purine ratio to resensitize these tumors to ICI (Keshet, Lee et al. 2020). In this context, our work has confirmed the key role of ASS1 in the resistance to ICI. Indeed, our data indicate that ASS1 is upregulated during the development of resistance to ICI and consequently could be used as a predictive biomarker. In addition, we have demonstrated that the intrinsic production of arginine through ASS1 and ASL is sufficient to impact mTORC1 and downstream mRNA translation. Interestingly, the targeting of ASS1 seems to have a dual effect on mRNA translation. On one side, inhibition of ASS1 downregulated eIF4F complex assembly through the inhibition of the phosphorylation of 4EBP1, and, on another side, favor the translation of a restricted subset of mRNA. This central role of mRNA translation control in the regulation of the sensitivity or the resistance of tumors to therapy is align with previous reports positing that mRNA translation reprogramming as central players of anti-tumor immune response (Falletta, Sanchez-Del-Campo et al. 2017, Cerezo, Guemiri et al. 2018, Han, Ji et al. 2019, Xu, Poggio et al. 2019, Suresh, Chen et al. 2020, Bartok, Pataskar et al. 2021, Zhang, Shi et al. 2025). In addition, a recent report highlighted the potential immunoregulatory role of ASS1 in ovarian cancer (Feng, Ji et al. 2022). Thus, our data can explain this first observation by describing a previously unknown role in translation regulation of antitumor immune response of ASS1. This discovery unravels a new actor in mRNA translation control, opening alternative windows to inhibit translation-driven plasticity of cancer cells and resistance to therapies.

In our experiments, the targeting of the enzymatic activity of ASS1 with MDLA, an analog of aspartate that induces a blockade of ASS1, recapitulates the effect of the genetic downregulation of ASS1. These results demonstrate that is the ASS1 enzymatic activity and consequently intracellular arginine concentration that is important in the regulation of mRNA translation controlling anti-tumor immunity and not an eventual moonlighting function like the ones that were described for other metabolic enzymes like GAPDH (Chang, Curtis et al. 2013).

These findings are significant for several reasons.

First, they demonstrate unexpected links between arginine synthesis pathways, mRNA translation control, and anti-tumor immunity. Arginine synthesis pathway on one side in part of the urea cycle is a crucial metabolite to sustain tumor cell proliferation as well as anti-tumor immunity. In the tumor-microenvironment and especially in melanoma, arginine is depleted in the tumor interstitial fluid compared to the plasma (Sullivan, Danai et al. 2019, Vecchio, Caiazza et al. 2021). This set up a competition for arginine access between cancer and stromal cells. This crucial needs of arginine were also the basement of the development of therapeutics approaches aiming to limit the arginine in the tumor microenvironment with for example the arginine deaminase (Sugimura, Ohno et al. 1992, Savaraj, Wu et al. 2007, Zhang, Liu et al. 2015). However, this type of strategy was not successful in the clinic with an absence of efficiency (Yao, Janku et al. 2021). In this context, our data demonstrating that ASS1 is upregulated during resistance to ICI suggest that it could be more beneficial to inhibit the arginine synthesis capacity of the cancer cells and through their plasticity than limiting the availability or arginine in the microenvironment. Indeed, previous report has highlighted the capacity of cancer cell to express ASS1 in low arginine condition to adapt and continue to proliferate while for CD8 T cells cannot and are fully dependent of the arginine import (Crump, Hadjinicolaou et al. 2021).

The second reason why our findings are noteworthy, is that we identify a new layer of regulation of mRNA translation through ASS1. Many studies have described the importance of mRNA translation in the context of antitumor immune response (Cerezo, Robert et al. 2021, Fabbri, Chakraborty et al. 2021). However, due to its pleiotropic role in the regulation of cell homeostasis in particular in immune cells, the direct targeting of translation could be detrimental for cancer treatment (Bjur, Larsson et al. 2013, Davari, Lichti et al. 2017). Our data indicated that ASS1 regulate mRNA translation demonstrate that the targeting of arginine synthesis and especially enzymatic activity of ASS1 can be used as an alternative to target mRNA translation reprogramming driving cancer aggressivity and resistance to therapies.

Finally, by demonstrating for the first time that ASS1 regulates mRNA translation and resistance to ICI,our work also positioned ASS1 and arginine synthesis pathway as potential biomarker. Even if further studies will be needed on large cohorts of melanoma patient to determine its theragnostic potential, over-expression of ASS1 and/or other enzyme from arginine synthesis pathway could be of remarkable therapeutic interest by allowing an early detection of ICI resistance events.

Altogether, the relationship that we have identified between arginine synthesis pathway and its limiting enzyme ASS1 with the mRNA translation initiation control and resistance to ICI highlights the need for a prompt development of drugs targeting metabolic and translational plasticity capacities of tumor cells with parallel translational studies in order to improve responses to cancer treatments and particularly ICI.

## Acknowledgements

We thank Mehdi Khaled for the YUMMs cell lines, Stephan Vagner for the pR–HepC–L bicistronic plasmid and Alexandre Puissant for the lentivirus packaging system. We thank the animal facility, the imaging facility, and the cytometry facility of C3M. We thank Jean-Erland Ricci, Stephan Vagner, Mehdi Khaled, Alexandre David, Mélanie Tichet as well as David M. Sabatini for helpful discussions. This study was possible due to the financial support from Fondation SILAB Jean Paufique, Fondation de France, Fondation ARC, Canceropôle SUD Provence-Alpes-Côte d’Azur, INCA, region SUD, INSERM First Step program and ANR JCJC TRANSMET.

## Author contributions

E.C. and W.K. design performed the experiments with the help of A.C.B.S.A.S., P.A. and G.B.; S.A. and S.R1. provided clinical samples; T.P. provide clinical samples and gave advices; I.B.S. team’s performed metabolomics experiments and gave advices; H.M. provide clinical samples and gave advices; C.R. gave access to polysome profiling platform, provide clinical samples and gave advices; S.S. gave advices and co-analyzed sequencing data with W.K.; E.C. S.R. and M.C. wrote the manuscript; S.R. and M.C. supervised all research and are joint senior authors.

## Methods

### Cell lines and reagents

The YUMM2.1, YUMM1.1, YUMM1.7 (murine melanoma) cell lines used in this study were a gift from Dr. M. Khaled. WM9 (human melanoma) cell line was obtained from ATCC. Cancer cell lines were maintained at 37°C and 5% CO_2_ in a humidified atmosphere and grown in DMEM medium supplemented with 10% fetal calf serum and penicillin/streptomycin (100U/mL) (Gibco). All cell lines were regularly verified to be mycoplasma-free using a mycoplasma detection assay, exploiting the activity of specific mycoplasmal enzymes that convert ADP to ATP. ATP is then detected via a bioluminescence signal (Lonza). YUMM1.1 and YUMM1.7 knocked-down for ASS1 were maintained in DMEM medium supplemented with L-argininosuccinic acid lithium salt 100µg.mL^-1^ (Sigma Aldrich). lZl-methyl-*DL*-aspartic acid (MDLA) was purchased from MedChem Express and was dissolved in PBS and the pH adjusted with NaOH to obtain a pH≈7.4.

### Metabolomic experiments

YUMM1.1 and YUMM2.1 cell lines were subjected to steady state metabolomic analyses accordingly to (Yuan, Kremer et al. 2019), using labelled ^15^N-amine-glutamine or ^15^N-amide-glutamine added to cell culture medium for 4 hours. Briefly, targeted tandem mass spectrometry (LC-MS/MS) was used to address the incorporation of non-radioactive stable isotope ^15^N into metabolites involved in the targeted pathways.

### Generation of stable knocked-down or over-expression cell lines

YUMM1.1, YUMM1.7 and YUMM2.1 cell lines were infected using lentiviral particles containing either shRNA targeting ASS1, scramble shRNA, non-regulatable form of ASS1 or an empty vector. The infected cells were selected using either puromycin or mCherry positive cell sorting analysis (Cytek Aurora Cell Sorter). These plasmids were purchased from Sigma-Aldrich or VectorBuilder and lentiviral particles produced in HEK293T cells using packing system vectors (psPAX2 and pCMV-VSVG plasmids) gifted by Dr. A. Puissant. ASS1 modulations were confirmed using western blot and qPCR analysis.

### Western-blot

Whole cell lysates were prepared in Ripa buffer containing protease and phosphatase inhibitors (Roche). The protein content in the cell lysates was quantified using DC Protein Assay (Biorad). Protein samples were resolved by SDS-PAGE, transferred onto a polyvinylidene fluoride (PVDF) membrane (Millipore). After saturation in a buffer containing BSA, gelatin, EDTA, Tween-20, TRIS-HCl and NaCl, membranes were then incubated with the appropriate antibodies. Proteins were visualized with the ECL System from Amersham.

### mRNA preparation, reverse transcription, and real time/quantitative PCR

Total RNAs were extracted using the RNeasy Minikit (Qiagen) following the manufacturer’s instructions. Reverse transcription was performed on 1*μ* g gof total RNA using the Reverse Transcription system kit (Promega) and random primers. Quantitative PCR was performed using Master Mix PowerUp SYBR Green (Applied Biosystems) on StepOne Thermocycler (Applied Biosytems). Melting curve analysis was used to verify the specificity of the generated amplicons. Relative mRNA levels were determined using the 2^-ΔΔCt^ method and *Actb* or *Tbp* as housekeeping genes.

### 35S-methionine labelling assay

Methionine incorporation experiments were performed according (Boussemart, Malka-Mahieu et al. 2014). Cells seeded in 6-well plates were washed 1 time with PBS and incubated 30 minutes in methionine free media. Cells were then incubated in a methionine-free media containing 11µCi.µL^-1^ of EasyTag EXPRESS ^35^S Protein Labeling Mix (PerkinElmer) for 30 minutes. For the cycloheximide (CHX) treated conditions, cells were incubated for 30 minutes with 100µg.mL^-1^ before methionine labelling and CHX was added in all the following steps. Cells were harvested in Ripa buffer containing protease and phosphatase inhibitors (Roche). The lysates were centrifuged at 12,000 rpm for 15 minutes and 20-30µL were spotted onto a Whatman 3mm filter paper. The proteins were precipitated with a cold 10% trichloroacetic acid (TCA) solution for 15 minutes followed by a 5% TCA solution for 10 minutes. The filter papers were then washed with 95% ethanol and air dried. The radioactivity was measured by scintillation, counting, and normalized protein content.

### pR-HeC-L-bicistronic dual luciferase reporter assays

The plasmid used in this assay was developed by Dr. Vagner and was used according to the methods described in (Cerezo, Guemiri et al. 2018). In this assay, translation of the first (LucR) cistron is eIF4F dependent, while translation of the hepatitis C virus internal ribosome entry site-driven second (LucF) cistron is eIF4F independent. Differential translation of both cistrons, as measured by the ratio of luciferase activities, directly reflects differences in eIF4F activity. Dual luciferase assays were conducted in indicated cell lines after transfection of 300ng of pR–HepC–L bicistronic vector in 24-well plates using jetPRIME transfection reagent (Polyplus). The lysates were prepared, and the luminescence measured in triplicate with the Dual-Luciferase Reporter Assay System, following the manufacturer’s instructions (Promega). The bioluminescence analyses shown are representative of at least 3 independent experiments.

### Proximity ligation assay

Two primary antibodies, raised in distinct species, are used to recognize two proteins of interest (e.g. eIF4E and eIF4G). Next, a pair of secondary antibodies (PLA probes) that are conjugated to complementary oligonucleotides and directed against the constant regions of each primary antibody (PLUS probe and MINUS probe) are used. Subsequent addition of a polymerase and HRP-labeled oligonucleotides result in an amplified rolling circular product. The signal resulting from each ligated pair of PLA probes is then visualized as an individual spot. These PLA signals can be quantified (counted) and assigned to a specific subcellular location based on microscopic images. The PLA protocol was followed according to the manufacturer’s instructions (Olink, Sigma-Aldrich). After blocking, the antibodies were used at the following concentrations: eIF4E (mouse, clone A-10, 1:200; SantaCruz Biotechnology, sc-271480) and eIF4G (rabbit, 1:200; Cell Signaling Technology, 2498) and incubated overnight with the primary antibodies at 4 °C. The PLA MINUS and PLA PLUS probes (containing secondary antibodies conjugated to oligonucleotides) were added and incubated for 1 h at 37°C. Ligase was used to anneal the two hybridized oligonucleotides into a closed circle. The DNA was then amplified (with rolling-circle amplification). Nuclei were stained with Olink mounting medium containing DAPI. The results were obtained with a confocal microscope (Nikon Confocal A1R) and the number of PLA signals per cell were counted (>3 fields) by semiautomated image analysis software (ImageJ and CellProfiler). The composite image was split into red and blue channels. The blue channel was used to detect and count the number of nuclei and the red channel to detect and quantify the number of dots. The results were reported as the ratio of dots to the number of detected cell nuclei.

### Polysomes profiling assay

Sucrose density gradient centrifugation was performed to separate the subpolysomal and polysomal ribosome fractions. Fifteen minutes prior to collection, the cells were incubated at 37 °C with 100 mg.ml^-1^ cycloheximide. Next, the cells were washed, scraped into ice-cold PBS supplemented with 100 mg.ml^-1^ cycloheximide, centrifuged at 3,000 r.p.m. for 5 min and then collected into 400 ml of LSB buffer (20 mM Tris, pH 7.4, 100 mM NaCl, 3 mM MgCl2, 0.5 M sucrose, 2.4% Triton X-100, 1 mM DTT, 100 U.ml^-1^ RNasin and 100 µg.ml-1 cycloheximide). After homogenization, 400 ml LSB buffer supplemented with 0.2% Triton X-100 and 0.25 M sucrose were added. The samples were centrifuged at 12,000g for 10 min at 4 °C. The resulting supernatant was adjusted with 5 M NaCl and 1 M MgCl2. The lysates were loaded onto a 15–50% sucrose density gradient and centrifuged in an SW41 rotor at 38,000 r.p.m. for 2 hours at 4 °C. The polysomal fractions were monitored and collected using a gradient fractionation system. Total RNA was extracted from the heaviest fractions and input samples using the TRIzol LS method.

### Translation efficiency analysis

Gene expression counts based on the exon alignments were used to statistically model the polysome profiling data using R software. Raw counts were first normalized using the TMM (trimmed mean of M-values) method implemented in edgeR to account for library size differences. Translation efficiency for each gene was calculated as the ratio of polysome-bound counts to total mRNA counts. Differential translation efficiency between experimental conditions was assessed by comparing the log2-transformed translation efficiency ratios across samples, and statistical significance of the observed differences was determined using edgeR.

Genes with a log_2_(FC) greater than 1.5 (*p* < 0.05) were considered significant genes, allowing us to identify the groups of genes upregulated at the translational level.

### *In vivo* experiments

Four-week-old immunocompetent male C57BL/6J mice were purchased from ENVIGO. All mice were Specific and Opportunistic Pathogen Free (SOPF). All procedures were conducted in accordance with French laws and European recommendations following a protocol approved by the French Ministry of Education and Research and by the local ethical committee for animal experimentation (CIEPAL Azur). All *in vivo* experimentations were performed in mice to acclimate for minimum one week in the animal facility. Mice were engrafted sub-cutaneously with 2x10^6^ cells in 100μL of PBS in the flank. When the tumor volumes reached a mean of 100mm^3^, mice were randomized and treated with the indicated compound. Antibodies *InVivo*Mab anti-mouse PD-1 (CD279), clone 29F.1A12 or *InVivo*Mab rat IgG2a isotype control, clone 2A3 were purchased from BioXCell, diluted in PBS and injected intraperitoneally (100µg in 100µL of PBS; 4mg.kg^-1^ per mouse). For the MDLA experiments, the same protocol was used. MDLA was injected intraperitoneally (14mg into 100µL of PBS + 10% of NaOH; 700mg.kg^-1^; pH 7.4), twice per day, every day, for 16 days. Anti-PD-1 or isotype IgG control treatment began 2 days after MDLA treatment. Tumors were measured every two days. Mice were sacrificed, all the same day, when one tumor reached 1000mm3, tumors were harvested for analysis by flow cytometry or immunofluorescence staining.

### Mice tumor dissociation

After mice were sacrificed, tumors were harvested and kept in RPMI media. Tumors were dissociated following the Tumor Dissociation Kit protocol (Miltenyi Biotec) and the dissociation using the gentleMACS Octo Dissociator (Miltenyi Biotec). Then, the cell suspensions were passed through a 70µm cell strainer, centrifugated at 400g for 5 minutes and stained as described in the flow cytometry assay method.

### Flow cytometry assay

After tumors dissociation, the cell pellets were incubated in FACS buffer (PBS, 1%BSA, 2mM EDTA) containing FcR blocking reagent (Miltenyi Biotec) for 30 minutes. After centrifugation cell pellets were incubated for 45 minutes with primary antibody, wash with PBS/EDTA/BSA containing 7AAD and then resuspended in 300µL PBS/EDTA/BSA. The stained cells were analyzed using a Spectral Aurora flow cytometer (Cytek). Datas were analyzed using FlowJo software.

### Immunofluorescence

Tumors were OCT embedded and snap frozen in liquid nitrogen. 5μm sections were obtained using a cryostat (Leica CM3050 S cryostat). Tissue sections were fixed using methanol (10 minutes at -20°) or PFA at 4% (10 minutes at room temperature). After 1hour of incubation with blocking buffer (PBS, BSA3%, normal goat serum 5%), primer antibodies were incubated overnight at 4° in PBS/BSA 1%. After 3 PBS washes, secondary antibodies were incubated for 1 hour at room temperature. Nuclei were stained with mounting media containing DAPI.

### Coimmunoprecipitation assay

For the co-immunoprecipitation experiment, WM9 melanoma cells were transfected with pRK5 CASTOR1-HA plasmid, obtained from Dr. D.M. Sabatini (Chantranupong, Scaria et al. 2016), and jetPRIME transfection reagent (Polyplus). Cells were maintained in DMEM and treated for 3 days with MDLA 10mM or cultivated for the last 12 hours in an arginine depleted media. Cells were immunoprecipitated using Pierce anti-HA magnetic beads (ThermoScientific) according to the manufacturer’s instructions. The immunoprecipitated complexes were analyzed by 10% SDSPAGE and immunoblotted using anti-HA antibody and anti-MIOS antibody.

### ASS1 enzymatic activity

For ASS1 enzyme activity, 1.0 × 10^6^ cells were seeded and treated with MDLA at the 0.5, 1, 5, or 10 mM for 72h. Cells were harvested using trypsin-EDTA 0.25%, washed with phosphate saline buffer, and cell pellets were obtained by centrifugation at 300 g for 5 min. Pellets were suspended in lysis buffer containing 20 mM Tris-HCl pH 7.2, 250 mM saccarose, 40 mM KCl, 2 mM EGTA, 1 mg/mL BSA plus protease and phosphatase inhibitors, and mechanical cell lysis was performed using a glass potter at 4 °C. Homogenates were centrifuged at 500 g for 20 min, supernatants were collected, pellets were resuspended in 1 mL fresh lysis buffer and centrifuged at 650 g for 20 min, and protein enriched homogenates were obtained by mixing both supernatants. Protein concentration was determined using the BCA assay (Thermo Fisher) and 5 µg protein were used per condition. Proteins were mixed with reaction buffer 20 mM Tris-HCl pH 7.8, 2 mM ATP, 2 mM citrulline, 2 mM aspartate, 6 mM MgCl_2_, 20 mM KCl, 0.2 U pyrophosphatase at 200 µL final volume. Then, the mixtures were incubated for 30 min at 37 °C and reaction was stopped by adding 200 µL stopping buffer (10 mM ascorbate, 2.5 mM ammonium molybdate and 2 % H_2_SO_4_). The ASS1 activities were measured spectrophotometrically at 650 nm in three technical replicates and minimum three biological replicates.

### Arginine dosage

Melanoma cells were plated in 6 wells plates, treated as indicated with MDLA for 72h and lysate in Ripa buffer. Arginine dosage was performed according to manufacturer instructions (Abcam ab241028).

### Patient-derived melanoma cell lines

Patient melanoma cells were prepared from fresh biopsies after obtaining informed consent from the patients. Briefly, biopsy was dissected and digested for 1 to 2 hours with collagenase A (0.33 U/mL), dispase (0.85 U/mL), and DNase I (144 U/mL) with rapid shaking at 37°C. Large debris were removed by filtration through a 70-mm cell strainer. Melanoma cells were then cultured in RPMI1640 medium supplemented with 10% fetal calf serum and penicillin/streptomycin (100U/mL) (Gibco).

### Statistical analysis

For statistical analysis, we used GraphPad Prism software v.9. Figure legends specify the statistical analysis used and define mean +/-SEM. *p*-values ≤ 0.05 were considered statistically significant.

## Supplementary Figure legends

**Figure S1:**
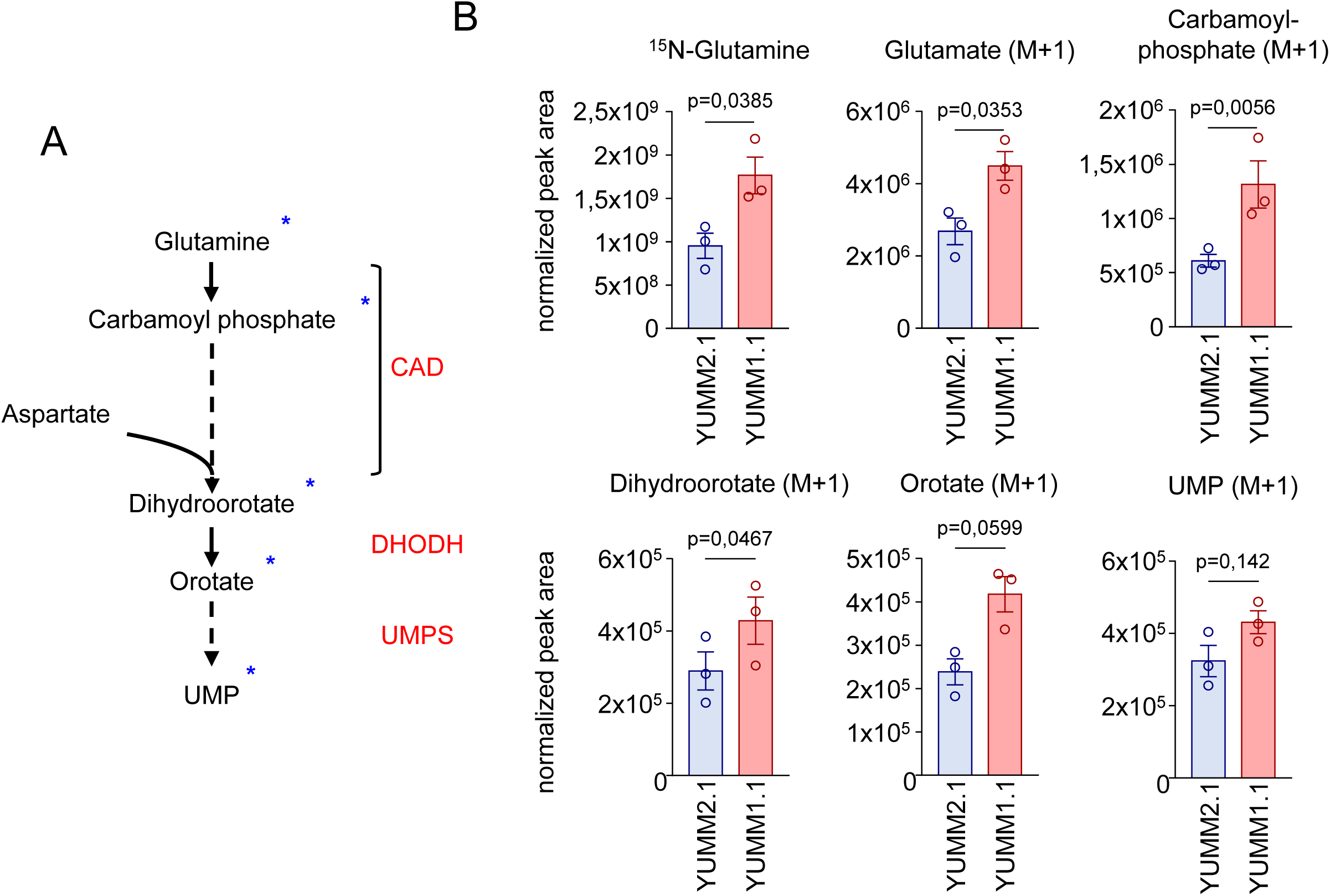
*De novo* pyrimidine synthesis is increase in ICI-resistant cells **A**, Schematic representation of the *de novo* pyrimidine synthesis fueled from glutamine. **D**, Normalized peak areas of ^15^N-labeled metabolites measured by targeted LC-MS/MS from YUMM2.1 sensitive and YUMM1.1 resistant melanoma cells.

**Figure S2:**
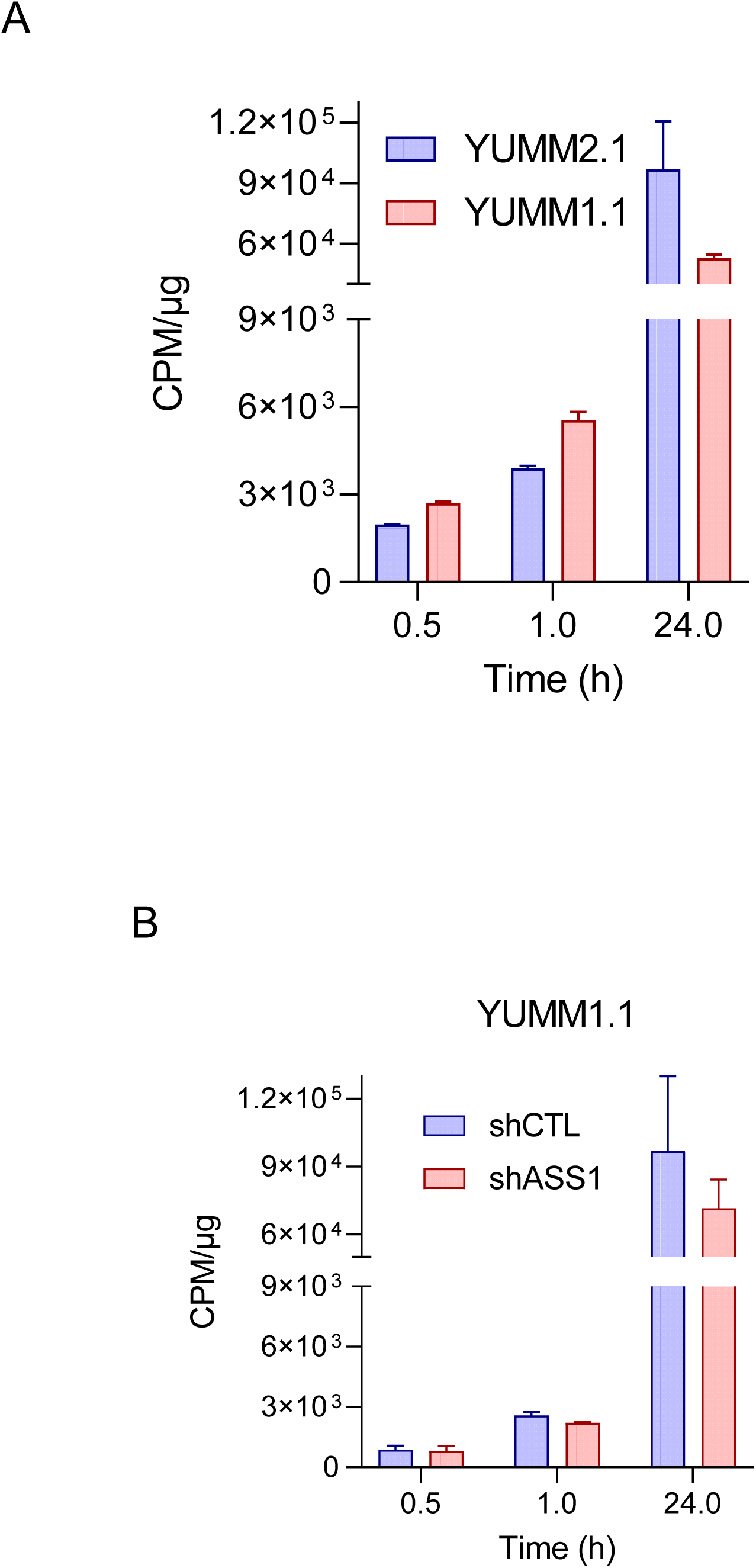
Modulation of ASS1 do not modulate arginine uptake **A**, [^3^H]-arginine uptake assays was performed on YUMM2.1 and YUMM1.1 for the indicated times. **B**, [^3^H]-arginine uptake assays was performed on YUMM1.1 expressing a shRNA targeting ASS1 or a shRNA control for the indicated times.

**Figure S3:**
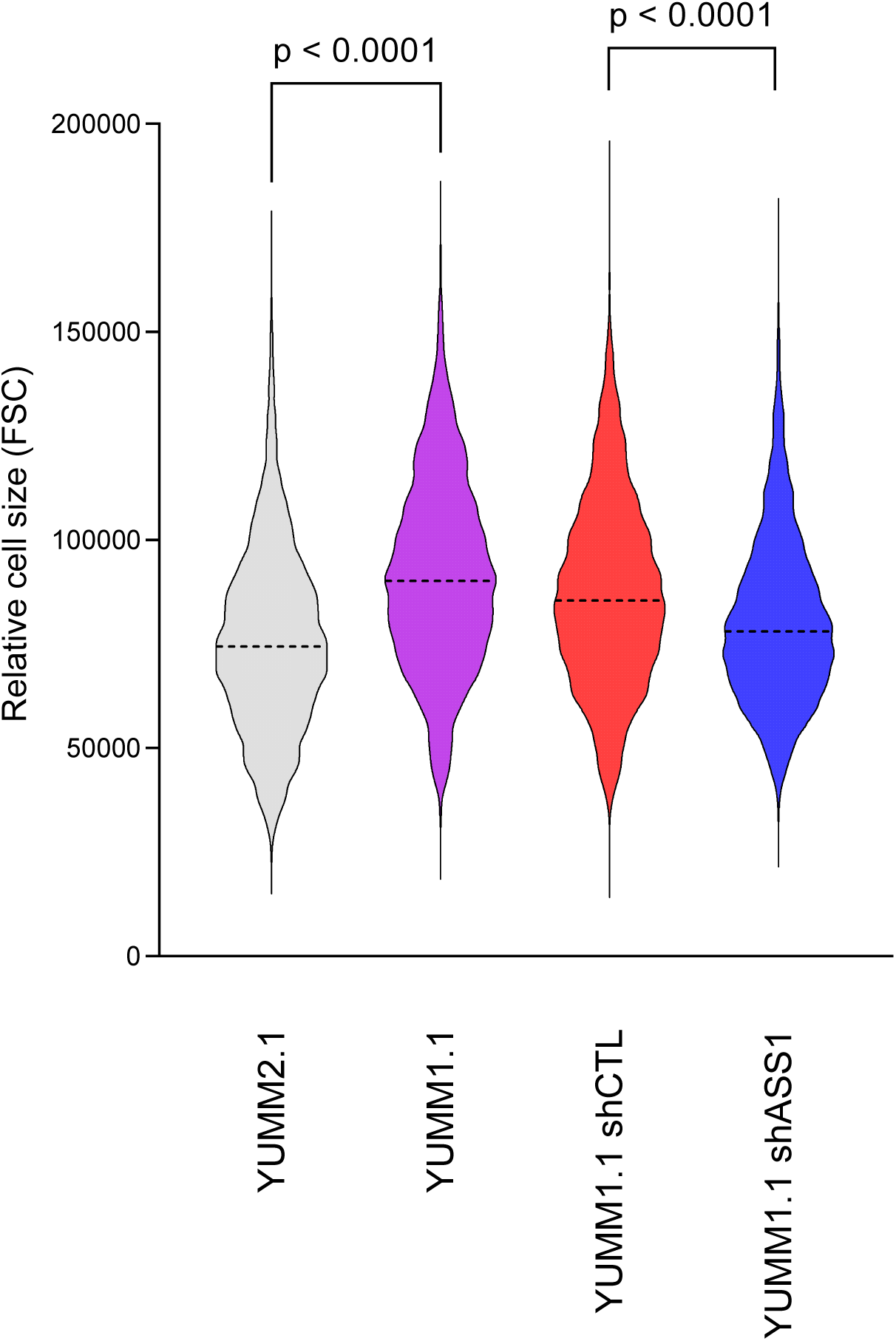
Modulation of ASS1 impact cell size The cell size of YUMM2.1, YUMM1.1 and YUMM1.1 expressing a shRNA targeting ASS1 or a shRNA control was estimated using flow cytometry.

**Figure S4:**
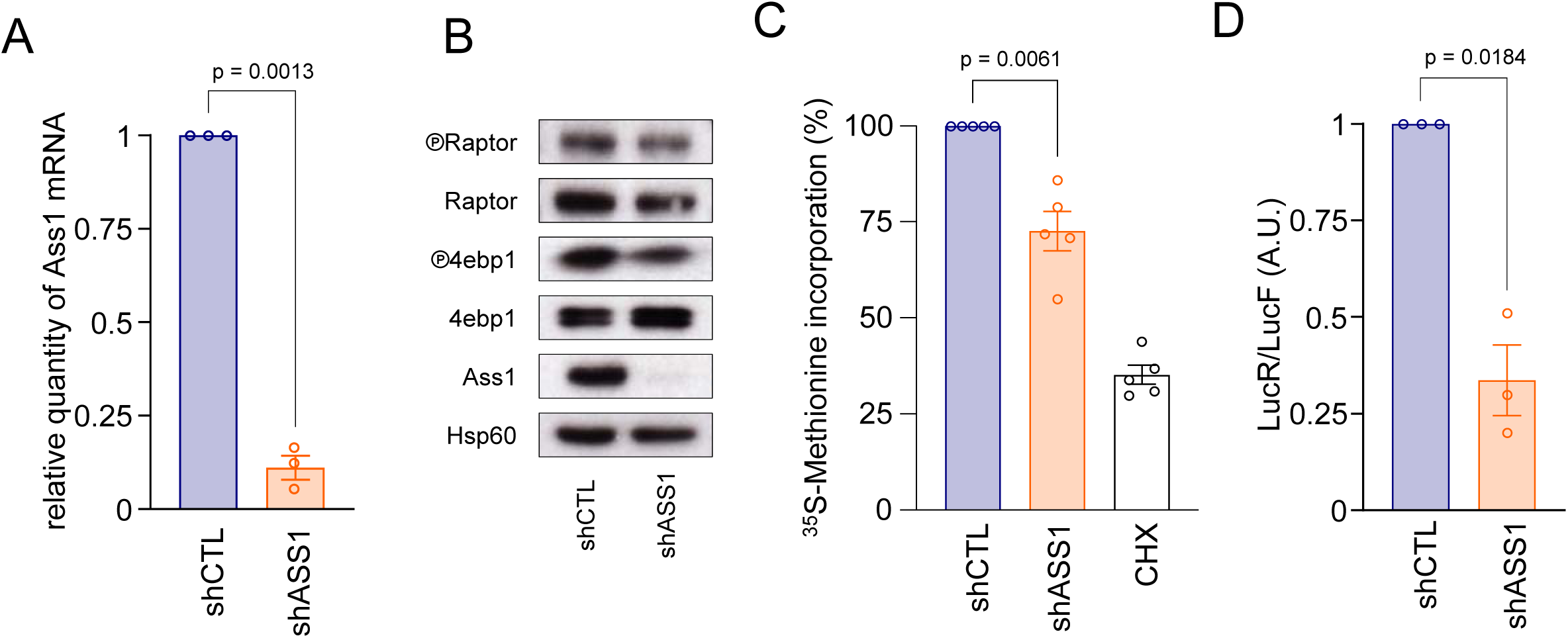
ASS1 inhibition decrease CAP-dependant translation **A**, RT-qPCR showing the expression level of ASS1 mRNA in YUMM1.1 expressing a second shRNA targeting ASS1 or a shRNA control. **B**, Western blot showing indicated proteins expression levels in YUMM1.1 cells expressing a second shRNA targeting ASS1 or a shRNA control. HSP60 serves as a loading control. **C**, Protein synthesis rates were determined in YUMM1.1 cells expressing a second shRNA targeting ASS1 or a shRNA control. Cells were then pulsed for 30 min with [^35^S]-Cys/Met, and the incorporation of [^35^S] into proteins was quantified and normalized to the total protein amount. Cells treated 30 minutes with 100µg/mL of cycloheximide (CHX) are used as a positive control. The data are presented as the mean (n = 5 ± SEM). **D**, Renilla over Firefly luminescent ratio quantification in YUMM1.1 cells expressing a second shRNA targeting ASS1 or a shRNA control (n=3; mean ± SEM).

